# Stability of spectral estimates in resting-state magnetoencephalography: recommendations for minimal data duration with neuroanatomical specificity

**DOI:** 10.1101/2021.08.31.458384

**Authors:** Alex I. Wiesman, Jason Da Silva Castanheira, Sylvain Baillet

## Abstract

The principle of resting-state paradigms is appealing and practical for collecting data from impaired patients and special populations, especially if data collection times can be minimized. To achieve this goal, researchers need to ensure estimated signal features of interest are robust. In electro- and magnetoencephalography (EEG, MEG) we are not aware of studies of the minimal length of recording required to yield a robust one-session snapshot of the frequency-spectrum derivatives that are typically used to characterize the complex dynamics of the brain’s resting-state. We aimed to fill this knowledge gap by studying the stability of common spectral measures of resting-state MEG source time series obtained from large samples of single-session recordings from shared data repositories featuring different recording conditions and instrument technologies (OMEGA: N = 107; Cam-CAN: N = 50). We discovered that the rhythmic and arrhythmic spectral properties of intrinsic brain activity can be robustly estimated in most cortical regions when derived from relatively short recordings of 30-s to 120-s of resting-state data, regardless of instrument technology and resting-state paradigm. Using an adapted leave-one-out approach and Bayesian analysis, we also provide evidence that the stability of spectral features over time is unaffected by age, sex, handedness, and general cognitive function. In summary, short MEG sessions are sufficient to yield robust estimates of frequency-defined brain activity during resting-state. This study may help guide future empirical designs in the field, particularly when recording times need to be minimized, such as with patient or special populations.

## 1. Introduction

The study of intrinsic brain activity during the resting-state is advancing our understanding of neural processes underlying a wide spectrum of brain functions in health and disease. Though the full functional relevance of the approach is still debated (Cole et al., 2010; Gonzalez-Castillo et al., 2021; Stevens and Spreng, 2014), the potential benefits of task-free paradigms are multifold. In electro- and magnetoencephalography (EEG, MEG), resting-state protocols lend themselves to a great variety of analysis approaches based on spectral transformations, including sophisticated derivatives of cross-frequency interactions (Canolty and Knight, 2010; Florin and Baillet, 2015) and functional connectivity (Hutchison et al., 2013). In principle, the fast temporal resolution of EEG and MEG may enable the estimation of such metrics across time, possibly over short, sliding time windows. A companion idea is to determine core individual features of brain activity in single individuals from the minimum possible amount of data, i.e., recording duration, especially in pediatric populations (Kurz et al., 2014; Kurz et al., 2018; Raschle et al., 2012), those with chronic pain (Davis et al., 2017; Diers et al., 2020; Witjes et al., 2021), and patients with cognitive impairment (Binnewijzend et al., 2012; Cassani et al., 2018; Mandal et al., 2018; Rombouts et al., 2005; Wiesman et al., 2021a; Wiesman et al., 2021b; Wiesman et al., 2018). Both aspects require the rigorous determination of brain signal stationarity and robustness of feature extraction.

Intra-session variability in resting-state activity has been of recent interest for “fingerprinting” (da Silva Castanheira et al., 2021; Finn et al., 2015), functional parcellation (Laumann et al., 2015), and enhanced prediction of behavior (Waschke et al., 2021) at the level of individual participants. Notwithstanding these recent efforts, most resting-state research derives and interprets summary statistics of neural fluctuations over data durations that vary considerably between studies, ranging at least from one (Ranasinghe et al., 2020) to fifteen (Marquetand et al., 2019) minutes. We are not aware of systematic studies of the within-session robustness of commonly used estimates of resting-state MEG signal features, nor of any resulting guidelines for minimum recording durations. Such guidelines would contribute to the evidence-based design of resting-state MEG studies that are more likely to be reproducible and would help researchers make informed decisions concerning key data collection parameters of shared repositories of MEG resting-state data.

Previous electrophysiological reliability studies have focused on the consistency of various signal metrics across sessions separated by days, months, or years (Duan et al., 2021; Fingelkurts et al., 2006; Garcés et al., 2016; Gasser et al., 1985; Keil et al., 2003; Lew et al., 2021; Liuzzi et al., 2017; Marquetand et al., 2019; McCusker et al., 2021; Thatcher, 2010). Here, we instead investigate the minimum recording duration required to achieve robust estimates of the main spectral properties of MEG resting-state source signals *within a single recording session*. Thus, we define a robust estimate as a signal derivative that exhibits stable, consistent values when computed from a sufficiently long duration of data recorded during a single session.

We also emphasize the recent interest in discriminating between periodic (i.e., rhythmic) and aperiodic (i.e., arrhythmic) components of the power spectrum of neurophysiological brain signals (Donoghue et al., 2020b). Studies have shown that their respective parameterizations help clarify the possible confounding contribution of background arhythmic brain activity to the measurement of oscillatory components of neural activity (Cellier et al., 2021; Donoghue et al., 2020a; Helfrich et al., 2021; Ostlund et al., 2021; Pani et al., 2021; Schaworonkow and Voytek, 2021; Van Heumen et al., 2021; Wilkinson and Nelson, 2021). Pathania et al. (2021) found that estimates of aperiodic components are consistent over intervals of at least seven days. We wish to expand these investigations by determining the minimum data duration required to produce robust estimates of the periodic and aperiodic components of MEG resting-state source maps.

Here we qualify an estimate obtained from a given data length as robust when its group-wise consistency exceeds pre-defined thresholds across measurements. We use intra-class coefficient (ICC) statistics, with thresholds of ICC > .90 for excellent, ICC > .75 for good, and ICC > .50 for moderate levels of estimate stability (Koo and Li, 2016). We examine the stability of spectral features derived from varying lengths of source-imaged MEG resting-state data collected from two different models of MEG instruments and available from two different shared repositories: the Open MEG Archive (OMEGA; CTF instrument; N = 107; Niso et al., 2016) and the Cambridge Centre for Ageing and Neuroscience dataset (Cam-CAN; Elekta instrument; N = 50; Taylor et al., 2017). We use the Fitting Oscillations & One-Over-F toolbox (FOOOF; Donoghue et al., 2020b) to decompose the spectra from MEG source time series into periodic and aperiodic components and examine the robustness of these components over varying data durations. Finally, we leverage the balanced demographic distribution and detailed cognitive testing of the Cam-CAN dataset to investigate potential linear effects of key demographics (i.e., age, sex, and handedness) and cognitive function (as measured by the Addenbrooke’s Cognitive Examination – Revised; ACE-R; Mioshi et al., 2006) on the intra-session stability of MEG resting-state spectral features, derived from multiple tested data durations. From these empirical results, we recommend minimum data durations, contingent on spectral and neuroanatomical features of interest, to ensure the robustness of spectral estimates from MEG resting-state data.

## 2. Methods

### 2.1 Data & participants

We used subsets of data from two shared repositories: the Open MEG Archive (OMEGA; Niso et al., 2016) and the Cambridge Centre for Ageing and Neuroscience dataset (Cam-CAN; Shafto et al., 2014; Taylor et al., 2017). Both OMEGA and Cam-CAN include resting-state MEG recordings from healthy adults, alongside basic demographic data and T1 MRIs. Key differences in the resting-state acquisition parameters between the two datasets include: the MEG system used (OMEGA: 275-axial gradiometer CTF, Port Coquitlam, BC, Canada; Cam-CAN: 204-planar gradiometer & 102-magnetometer MEGIN/Elekta VectorView, Helsinki, Finland), the data acquisition site (OMEGA: Montreal, QC, CA; Cam-CAN: Cambridge, UK), basic participant behaviour (OMEGA: eyes-open; Cam-CAN: eyes-closed), and the sampling rate (OMEGA: 2400 Hz; Cam-CAN: 1000 Hz).

From the OMEGA dataset, 107 participants (mean age = 30.57 [SD = 12.70]; age range = 19 – 74 years; 102 right-handed; 46 female) were included based on the following criteria: no current neurological or psychiatric disorder, no history of head trauma, no MRI or MEG contraindications, and complete and useable MEG, MRI, and demographic data. Eyes-open resting-state MEG data were collected from each participant at a sampling rate of 2400 Hz and with an antialiasing filter with a 600 Hz cut-off. Noise-cancellation was applied using CTF’s built-in third-order spatial gradient noise filters. Recordings lasted a minimum of 4 minutes and were conducted with participants in the seated position as they fixated on a centrally-presented crosshair.

We also selected a validation sample of 50 healthy adult participants (mean age = 44.72 [SD = 14.56]; age range = 20 – 69 years; 45 right-handed; 23 female) from the Cam-CAN dataset. The sample size of 50 was determined quasi-empirically, by modeling the time-by-ICC stability relationship as a function of sample size in the OMEGA sample (see *2.4 Temporal stability analysis* and Figure 1B-2). From these relationships, it was apparent that a sample of at least 40-50 participants was necessary for the time-by-ICC estimation to stabilize, and for the variability in this estimation due to random participant subsampling to diminish (Figure S1). To study the impact of age on the stability of MEG derivatives, 10 participants were selected per each decade from 20 – 69 years of age, with the other demographics matched to those of the OMEGA participant sample. Exclusionary criteria included current neurological or psychiatric disorder, MRI or MEG contraindications, unusable MEG, MRI, or demographic data, and cognitive impairment (MMSE ≤ 24). Resting-state MEG data were collected from each participant at a sampling rate of 1000 Hz and with a band-pass filter of 0.03 – 330 Hz. Noise-cancellation was applied using tSSS/MaxFilter (v2.2; MEGIN/Elekta). Recordings lasted approximately 8 minutes with participants in the seated position and their eyes closed.

**Figure 1.**
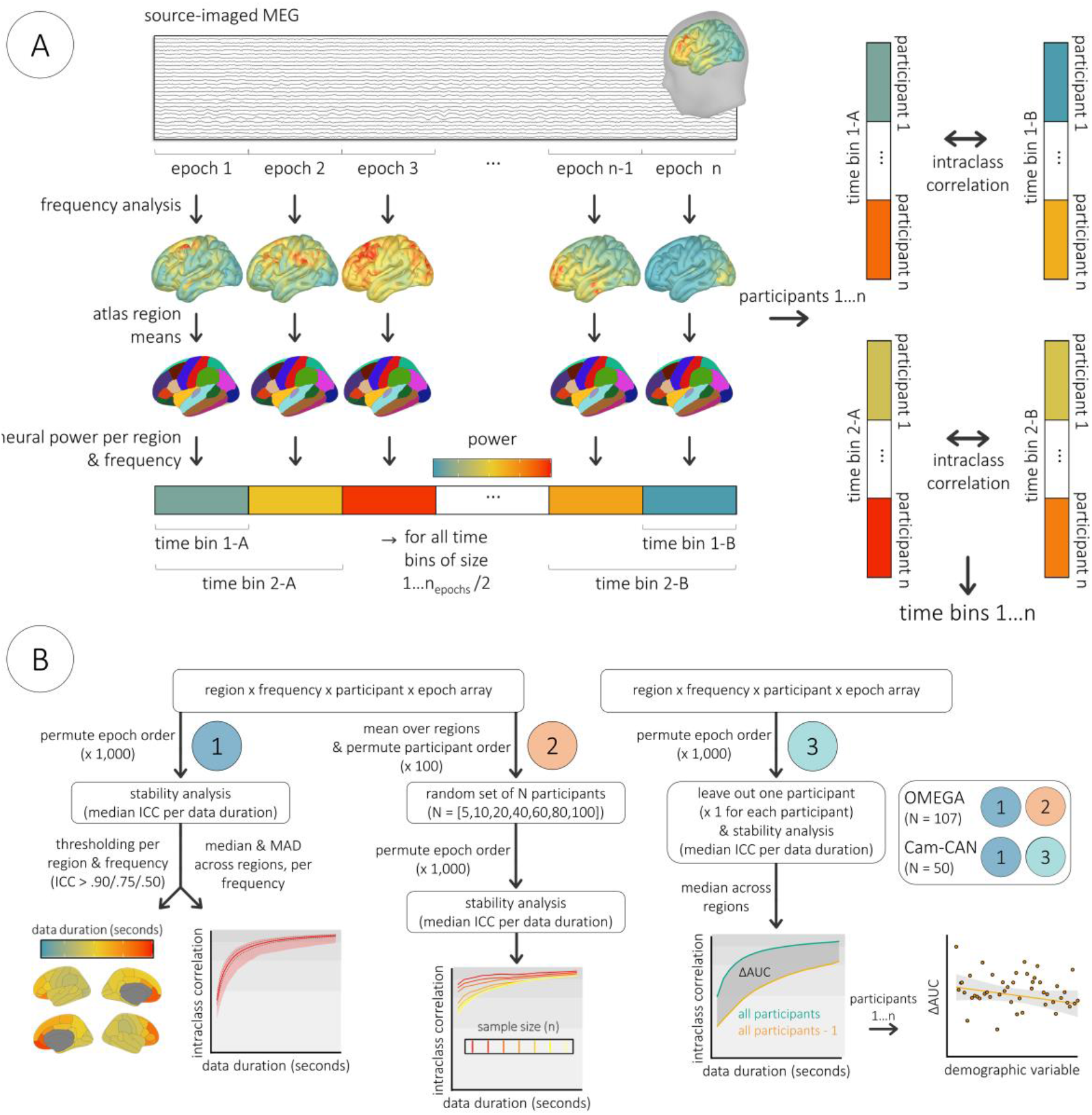
Stability analysis pipeline. Source-imaged MEG data was epoched into consecutive, non-overlapping segments and transformed into the frequency domain per vertex. Power spectral density values were then averaged within canonical frequency bands and over vertices within Desikan-Killiany atlas regions. For each of 1,000 permutations, the temporal order of these epochs was randomized. Data for each cortical location, frequency, and participant were then averaged over time bins of progressively larger numbers of epochs from one to n_epochs_/2 and used to compute ICC with time bins averaged over the same number of different epochs. For instance, within each permutation for participants 1…n, spectral power estimates from the first epoch (time bin 1-A) were correlated with spectral power estimates derived from the last epoch (time bin 1-B), spectral power estimates averaged over the first two epochs (time bin 2-A) were then correlated with spectral power estimates averaged over the last two epochs (time bin 2-B), and so forth until all epochs were included (two time bins each averaged over n_epochs_/2 epochs, i.e., 1/2 of the total recording duration). Computing the median across these permutations for each size of time bin (from 1 to n_epochs_/2) resulted in a time course of intra-session ICC values per each combination of region and frequency. (B) Graphical workflows. All analysis steps are indicated for the derivation of (1) stability estimates per each combination of region and frequency, (2) stability estimates per each combination of frequency and sample size, and (3) change scores representing the omission of each participant from the stability analysis. The extra inset to the far right indicates the participant subsamples on which each workflow was implemented. AUC: area under the curve. Cam-CAN: Cambridge Centre for Ageing and Neuroscience participant subsample. ICC: intraclass correlation coefficient. MAD: median absolute deviation. OMEGA: Open MEG Archives participant subsample.

The data collection and management protocols for the OMEGA and Cam-CAN repositories were approved by the research ethics boards at the Montreal Neurological Institute and the University of Cambridge, respectively. All participants in both studies provided written informed consent in accordance with the Declaration of Helsinki.

### 2.2 MEG data preprocessing

MEG preprocessing procedures for both datasets were similar to those reported by da Silva Castanheira et al. (2021), following good-practice guidelines (Gross et al., 2013). All processing steps were performed using Brainstorm (Tadel et al., 2011), unless otherwise specified. Notch filters were applied at the respective line-in frequency (and harmonics) of each dataset (60 Hz for OMEGA, 50 Hz for Cam-CAN), along with a 0.3 Hz high-pass FIR filter to attenuate slow-wave drift and DC offset. Additional notch filters were applied at 88 Hz (and harmonics) to attenuate known artifacts in the Cam-CAN dataset. Signal space projectors (SSPs) were derived around cardiac and eye-blink events detected from ECG and EOG channels and applied to the data. SSPs were also used to attenuate low-frequency (1 – 7 Hz) and high-frequency (40 – 400 Hz) noise related to saccades and muscle activity.

### 2.3 Source imaging & spectral analysis

Using approximately 100 digitized head points, MEG data were co-registered to each participant’s T1-weighted MRI that was segmented and labeled with Freesurfer (Fischl, 2012) using *recon-all*. Source imaging was performed using only gradiometer data with individually fitted overlapping-spheres forward models (15,000 vertices, with elementary sources constrained normal to the cortical surface) and Brainstorm’s linearly-constrained minimum variance (LCMV) beamformer with default parameters, and noise covariance estimated from empty-room recordings taken as close in time as possible to each participant recording. This source imaging procedure yielded continuous time series per each spatially-resolved location (i.e., vertex) on the cortical surface for each participant.

Source-imaged data were segmented into contiguous, non-overlapping 6-second epochs, resulting in a total of 40 epochs (total of 240 seconds) for OMEGA participants and 70 epochs (total of 420 seconds) for Cam-CAN participants. Vertex-wise estimates of power spectral density (PSD) were obtained using Welch’s method (3-second time window, 50% overlap) and averaged over canonical frequency bands using Brainstorm defaults (delta: 2 – 4 Hz; theta: 5 – 7 Hz; alpha: 8 – 12 Hz; beta: 15 – 29 Hz; low-gamma: 30 – 59 Hz; high-gamma: 60 – 90 Hz). To examine the potential effect of frequency bandwidth on subsequent analyses, a complementary procedure was performed in the OMEGA participants across the same approximate frequency ranges using constant spectral increments (i.e., from 3 – 92 Hz in 10-Hz steps). To study the effects of epoch/window length on spectral parametrization (see *2.5 Stability of power spectrum parameterization*), we also estimated PSDs from 12-second epochs using 6-second time windows in the OMEGA sample (also with 50% overlap). Within each frequency band and for each participant, PSD values were averaged over vertices belonging to each of the 68 regions of a standard atlas (Desikan et al., 2006) registered to individual structural data (Freesurfer). This resulted in an array of PSD values of number of atlas regions x number of frequency bins x number of participants x number of epochs (e.g., 68 × 6 × 107 × 40 from OMEGA).

### 2.4 Temporal stability analysis

The stability of spectral estimates was assessed via intraclass correlation coefficients (ICC; single-rater, two-way mixed-effects, absolute agreement; Koo and Li, 2016; McGraw and Wong, 1996) between multi-dimensional PSD arrays averaged across various numbers of randomly selected, non-overlapping epochs (Figure 1A). To control the potential bias that could be due to the epoch’s temporal position in the session, ICC derivations were repeated across 1,000 randomizations of the epoch order: for each of 1,000 permutations, the order of the matrix data was randomized over epochs using the *randperm* function in MATLAB (version 2019b; Mathworks, Inc., Massachusetts, USA), and ICCs were calculated for each region and frequency between sets of PSD estimates averaged over progressively larger numbers of epochs from one to n_epochs_/2 (using the *ICC.m* function in Matlab; Salarian, 2016). The medians of these values were then computed across randomizations, leading to estimates of intraclass similarity for each cortical region and spectral frequency as a function of recording length (Figure 1B-1). From these data, the minimum durations where ICC exceeded established thresholds (Koo and Li, 2016) for moderate (ICC > .50), good (ICC > .75), and excellent (ICC > .90) reliability were extracted for each combination of region with frequency and plotted on a spatial representation of the Desikan-Killiany atlas using *ggseg* (Mowinckel and Vidal-Piñeiro, 2020). Additionally, the median, median absolute deviation, and range of ICC values across regions were computed and plotted as a function of time and frequency using *ggplot2* (Wickham, 2011). To examine the potential effect of group size on stability, the original 4-dimensional PSD arrays were averaged across all regions and a similar approach was taken to test for the robustness of spectral estimates across various sample sizes of randomly selected participants (N = [5, 10, 20, 40, 60, 80, 100]; Figure 1B-2). One hundred iterations of the participant randomization were performed and within each of these randomizations the previously described stability estimation approach was followed (1,000 epoch permutations). Medians and median absolute deviations of the resulting time-by-ICC estimates were calculated across the 100 participant randomizations to examine their stabilization with increasing sample sizes.

### 2.5 Stability of power spectrum parameterization

We parameterized the power spectra of each brain region using the FOOOF algorithm (Donoghue et al., 2020b) implemented in Brainstorm (MATLAB version; frequency range = 0.5 – 40 Hz; Gaussian peak model; peak width limits = 0.5 – 12 Hz; maximum n_peaks_ = 3; minimum peak height = 3 dB; proximity threshold = 2 SD; fixed aperiodic; no guess weight). The robustness of estimated aperiodic features from the FOOOF models (i.e., the exponent and offset) over varying epoch lengths was tested using the procedure detailed above. In addition, the periodic (i.e., rhythmic) features were averaged over the same canonical frequency bands described earlier (see *2.3 Source imaging & spectral analysis*) and the robustness over varying epoch lengths tested using the above-mentioned pipeline. To examine whether the goodness of fit of FOOOF models might bias the stability of parameterized features, we regressed each feature on the model fit (i.e., R2) per each relevant location, frequency, and epoch (i.e., with participants as the unit of measurement; using the *fitglm* function in MATLAB) and extracted residuals from these linear models. We then re-ran all FOOOF temporal stability analyses on the fit-corrected FOOOF feature residuals.

### 2.6 Individual stability contribution analysis

To determine whether key demographic and cognitive factors biased the estimation of MEG temporal stability, we computed an *individual stability contribution* score per participant in the Cam-CAN participant sample (Figure 1B-3), and related these scores to age, sex, handedness, and general cognitive function (as measured by the ACE-R; Mioshi et al., 2006). We used a leave-one-out adaptation of the previously described stability analysis, wherein for each randomization, time-by-ICC models were generated using all but one participant. The median of these ICC values across brain regions was then extracted per time sample and frequency, leading to a single time-by-ICC model per frequency band. The area under the curve (AUC) was calculated for all such models using the *trapz* function in MATLAB. Each missing-participant’s stability model AUC value was subtracted from a full-sample model AUC value generated within the same set of epoch permutations. This resulted in a single ΔAUC score for each Cam-CAN participant per frequency, representing the change in the temporal stability of the model when they were excluded, relative to when using the full sample. The higher the value, the more stable the model when including said participant. These scores were individually regressed against demographic and cognitive data using the *lm* function in *R* (Team, 2017). Bayesian testing of these models was performed using the *BayesFactor* package (Morey et al., 2015).

### 2.7 Code & data availability

Data used in the preparation of this work were obtained from the Cam-CAN repository (available at http://www.mrc-cbu.cam.ac.uk/datasets/camcan/; Shafto et al., 2014; Taylor et al., 2017) and the OMEGA repository (available at https://www.mcgill.ca/bic/resources/omega; Niso et al., 2016). Code for MEG preprocessing and the stability analysis is available at https://github.com/aiwiesman/rsMEG_StabilityAnalysis.

## 3 Results

### 3.1 Temporal stability of intra-session MEG across time & space

The minimal recording durations required to derive robust, stable estimates of MEG spectral features for OMEGA (n = 107) and Cam-CAN (n = 50) are illustrated in Figures 2 & 3 and summarized in Tables 1 & 2. Despite substantial differences in sensing technology, recording environment, and participant demographics, the results were similar between both samples. Overall, most spectral features were robustly estimated when derived from 30 to 120 seconds of data. In the alpha (8 – 12 Hz) and beta (15 – 29 Hz) bands, activity in every region showed “excellent” stability (ICC > .90) when derived from data durations of 2 minutes or less, while activity in the delta (2 – 4 Hz), theta (5 – 7 Hz) and low-gamma (30 – 59 Hz) bands reached excellent stability in every region with less than 3.5 minutes of data. High-gamma (60 – 90 Hz) neural activity reached “good” stability in every region but one (left pars opercularis) with data durations of 3.5 minutes or less. Differences in stability across canonical frequency bands were not due to differences in bandwidth: the tests conducted with the OMEGA sample across equivalent bandwidths of 10 Hz produced similar stability results (Figure S2).

**Table 1.** Values are data durations (in seconds) required to reach specified levels of excellent, good, and moderate stability. Median, minimum, and maximum values were computed across brain regions, and the most and least stable regions were defined as the regions that exhibited the highest and lowest overall intraclass correlations, respectively, across all possible data durations. DNS: did not stabilize to specified ICC level within available data durations. ICC: intraclass correlation coefficient. OMEGA: Open MEG Archives participant subsample.

| OMEGA |  | excellent – ICC > .90 |  |  | good – ICC > .75 |  |  | moderate – ICC > .50 |  |  | most stable region | least stable region |
| --- | --- | --- | --- | --- | --- | --- | --- | --- | --- | --- | --- | --- |
|  |  | median | min | max | median | min | max | median | min | max |  |  |
| power spectral density | delta (2 - 4 Hz) | 72 | 42 | *DNS* | 24 | 12 | 60 | 6 | 6 | 18 | postcentral L | frontalpole L |
|  | theta (5 - 7 Hz) | 96 | 66 | *DNS* | 30 | 24 | 48 | 12 | 6 | 18 | precentral L | parahippocampal R |
|  | alpha (8 - 12 Hz) | 72 | 54 | 114 | 24 | 18 | 36 | 12 | 6 | 12 | parsorbitalis R | cuneus L |
|  | beta (15 - 29 Hz) | 33 | 18 | 60 | 12 | 6 | 18 | 6 | 6 | 6 | temporalpole L | supramarginal R |
|  | low-gamma (30 - 59 Hz) | 72 | 12 | *DNS* | 24 | 6 | *DNS* | 6 | 6 | 78 | superiorparietal L | parsopercularis R |
|  | high-gamma (60 - 90 Hz) | *DNS* | 42 | *DNS* | 111 | 12 | *DNS* | 36 | 6 | *DNS* | superiorparietal L | parstriangularis R |
| parameterized periodic | delta (2 - 4 Hz) | *DNS* | *DNS* | *DNS* | 114 | 48 | *DNS* | 39 | 18 | 72 | postcentral L | frontalpole L |
|  | theta (5 - 7 Hz) | 69 | 36 | *DNS* | 24 | 12 | 54 | 6 | 6 | 18 | lateraloccipital L | frontalpole L |
|  | alpha (8 - 12 Hz) | 42 | 30 | 72 | 12 | 12 | 24 | 6 | 6 | 12 | superiortemporal L | frontalpole L |
|  | beta (15 - 29 Hz) | 30 | 18 | 54 | 12 | 6 | 18 | 6 | 6 | 6 | precentral L | frontalpole R |
| parameterized aperiodic | slope | 36 | 24 | *DNS* | 12 | 6 | 42 | 6 | 6 | 18 | postcentral R | frontalpole R |
|  | offset | 45 | 30 | 120 | 18 | 12 | 42 | 6 | 6 | 12 | postcentral R | frontalpole L |

**Table 2.**
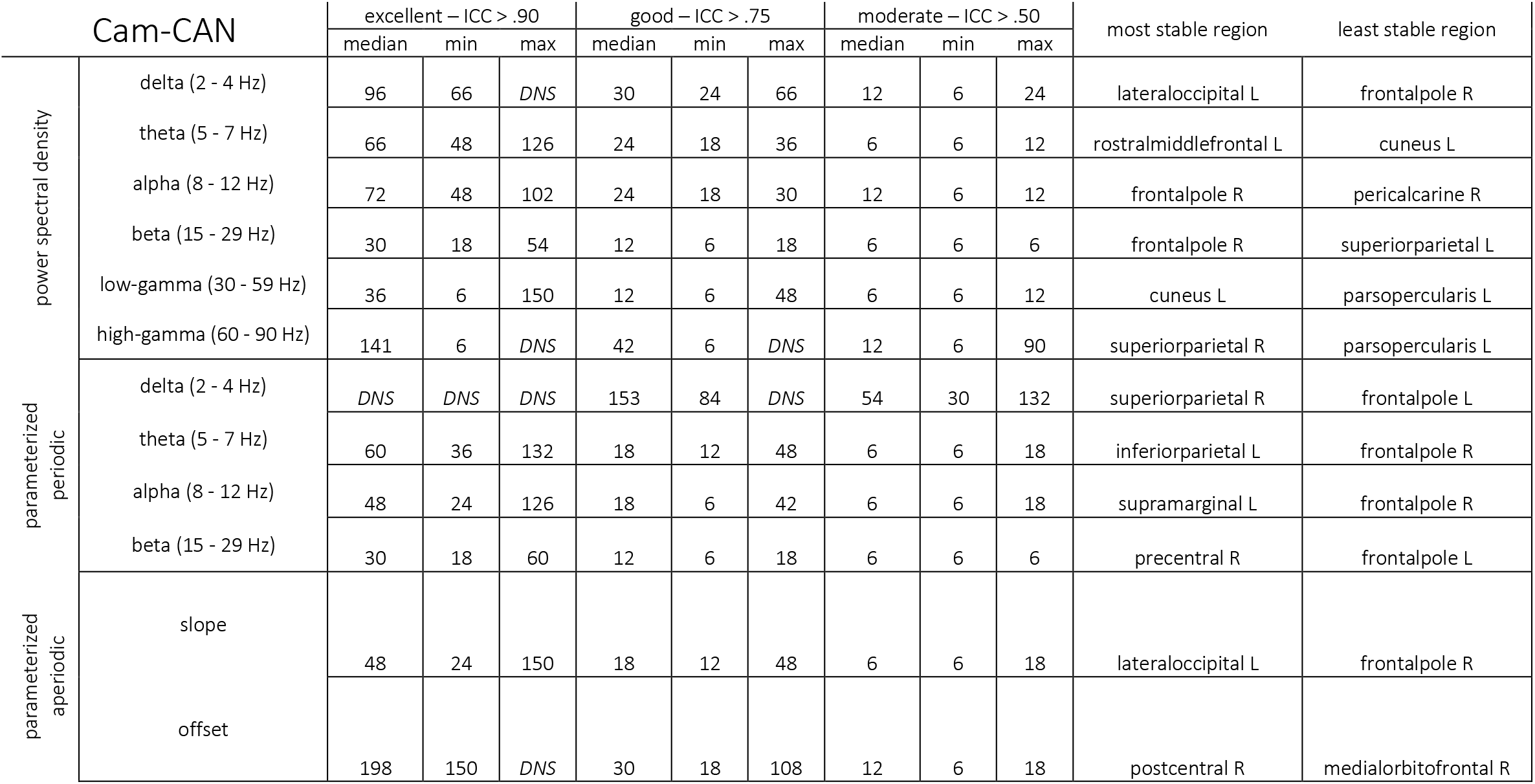
Values are data durations (in seconds) required to reach specified levels of excellent, good, and moderate stability. Median, minimum, and maximum values were computed across brain regions, and the most and least stable regions were defined as the regions that exhibited the highest and lowest overall intraclass correlations, respectively, across all possible data durations. DNS: did not stabilize to specified ICC level within available data durations. ICC: intraclass correlation coefficient. Cam-CAN: Cambridge Centre for Ageing and Neuroscience participant subsample.

**Figure 2.**
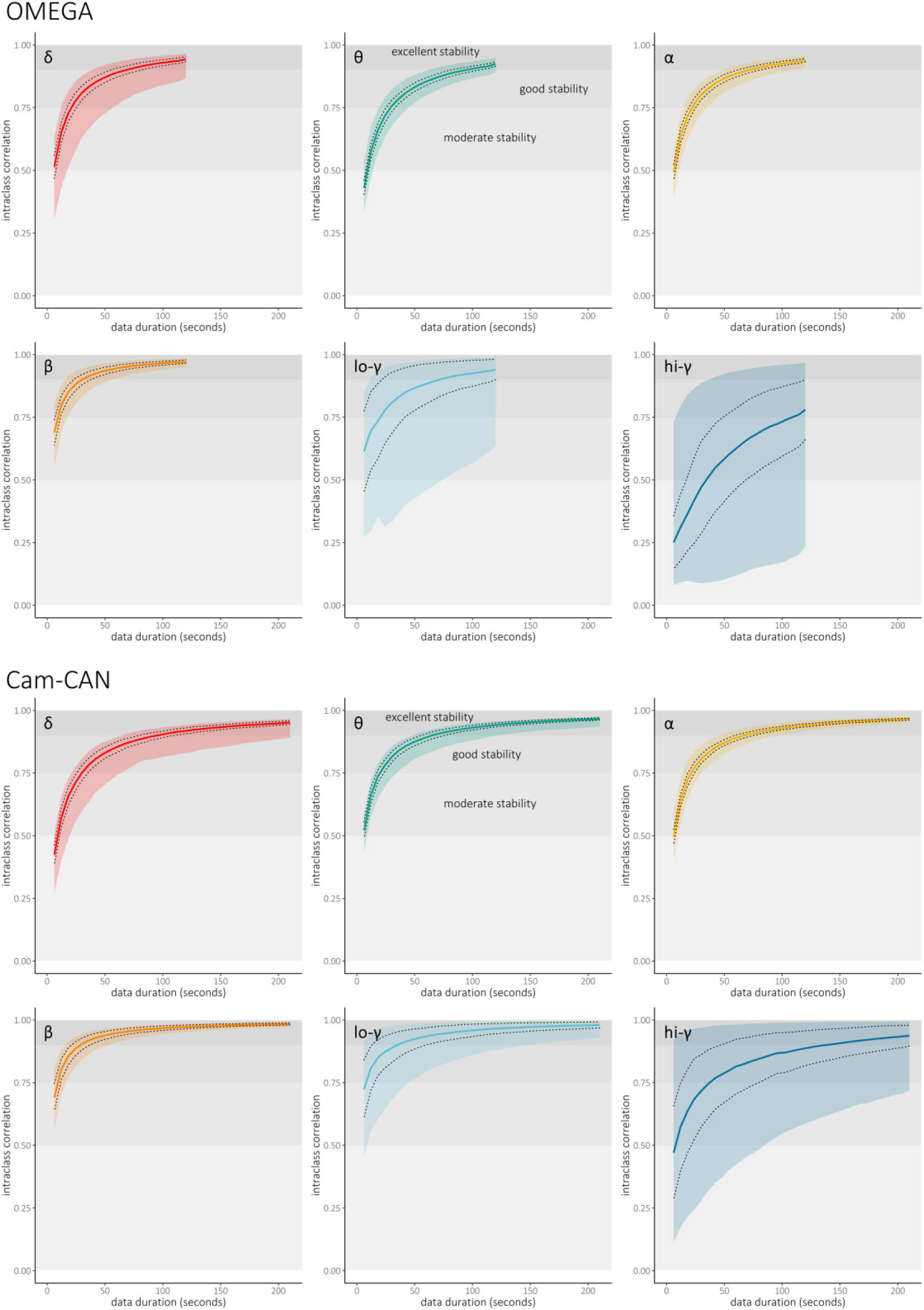
Intra-session temporal stability of band-limited power. Intraclass correlation coefficients (y-axes) were obtained from 1,000 permutations of epoch order and represent a measure of the stability of spectral power for each frequency band of interest as a function of data duration (x-axes, in seconds; top: OMEGA, N = 107; bottom: Cam-CAN, N = 50). Colored lines represent the median across regions, dotted lines indicate ± one median absolute deviation across regions, and colored shaded intervals represent the range of stability values across all modeled cortical regions of the Desikan-Killiany atlas. Horizontal shaded intervals in each plot represent typical thresholds used to define moderate (ICC > .50), good (ICC > .75), and excellent (ICC > .90) reliability. Note that the maximum data duration from OMEGA was shorter than from Cam-CAN (see Methods).

**Figure 3.**
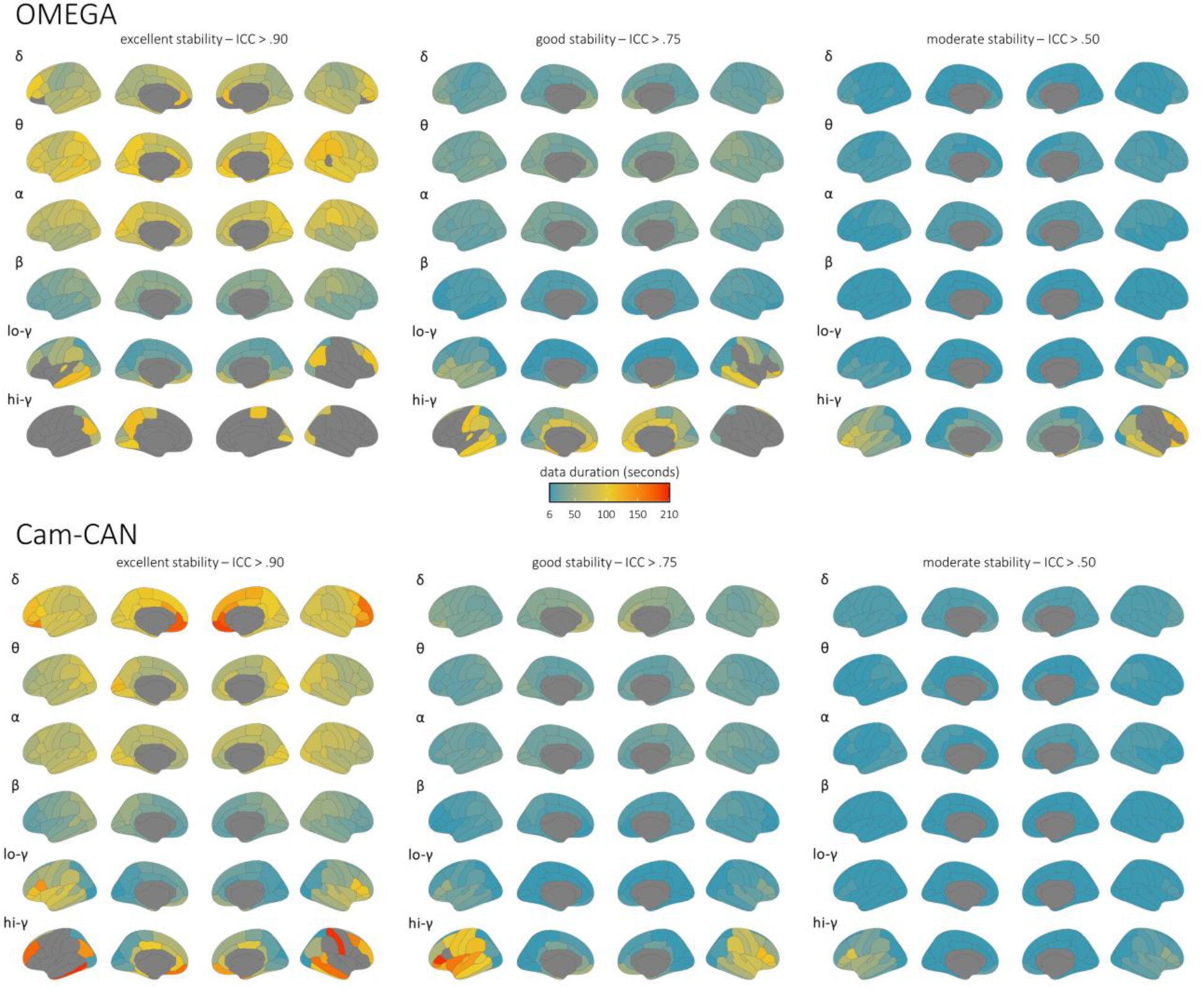
Brain maps of intra-session stability of spectral power estimates. Parcellated surface maps per each spectral frequency (denoted by Greek letters to the top-left of each set) indicate the length of data (color bar; in seconds) required to achieve accepted thresholds for moderate (ICC > .50; right), good (ICC > .75; middle), and excellent (ICC > .90; left) reliability in each participant sample (top: OMEGA, N = 107; bottom: Cam-CAN, N = 50) for each modeled cortical region of the Desikan-Killiany atlas. Warmer colors indicate worse temporal stability, and regions left grey did not achieve the indicated level of reliability within the maximum length of data available for the respective participant sample (OMEGA: 120 seconds; Cam-CAN: 210 seconds). Note that these are spatial representations of the same data shown in Figure 2.

### 3.2 Stability of aperiodic & periodic spectral components

In both the OMEGA and Cam-CAN samples, the estimation of the aperiodic slope achieved excellent stability when derived from less than 2 minutes of data. However, the estimation of the aperiodic offset was noticeably less robust in Cam-CAN participants, particularly in frontal cortices (Figure 4). Parameterized periodic neural activity in the theta, alpha, and beta bands reached excellent stability in every region with less than 2 minutes of data, apart from bilateral inferior and medial frontal regions (Figure 5), which required slightly longer data durations. In contrast, the periodic delta components were noticeably less stable, and did not reach excellent stability in any region with 3.5 minutes of data. Importantly, the instability of the delta periodic component was neither due to the length of the epoch nor of the time window used for PSD derivations: similar results were obtained when using 6-s epochs/3-s time windows or 12-s epochs/6-s time windows (Figures S3 & S4). The instability of the delta periodic component may instead have been caused by intra-session fluctuations in the goodness-of-fit of the FOOOF model at low frequencies. Controlling for regional variations in FOOOF model fit (i.e., R2) across participants in the OMEGA sample markedly improved the robust estimation of periodic activity in the delta band (Figures S5 & S6).

**Figure 4.**
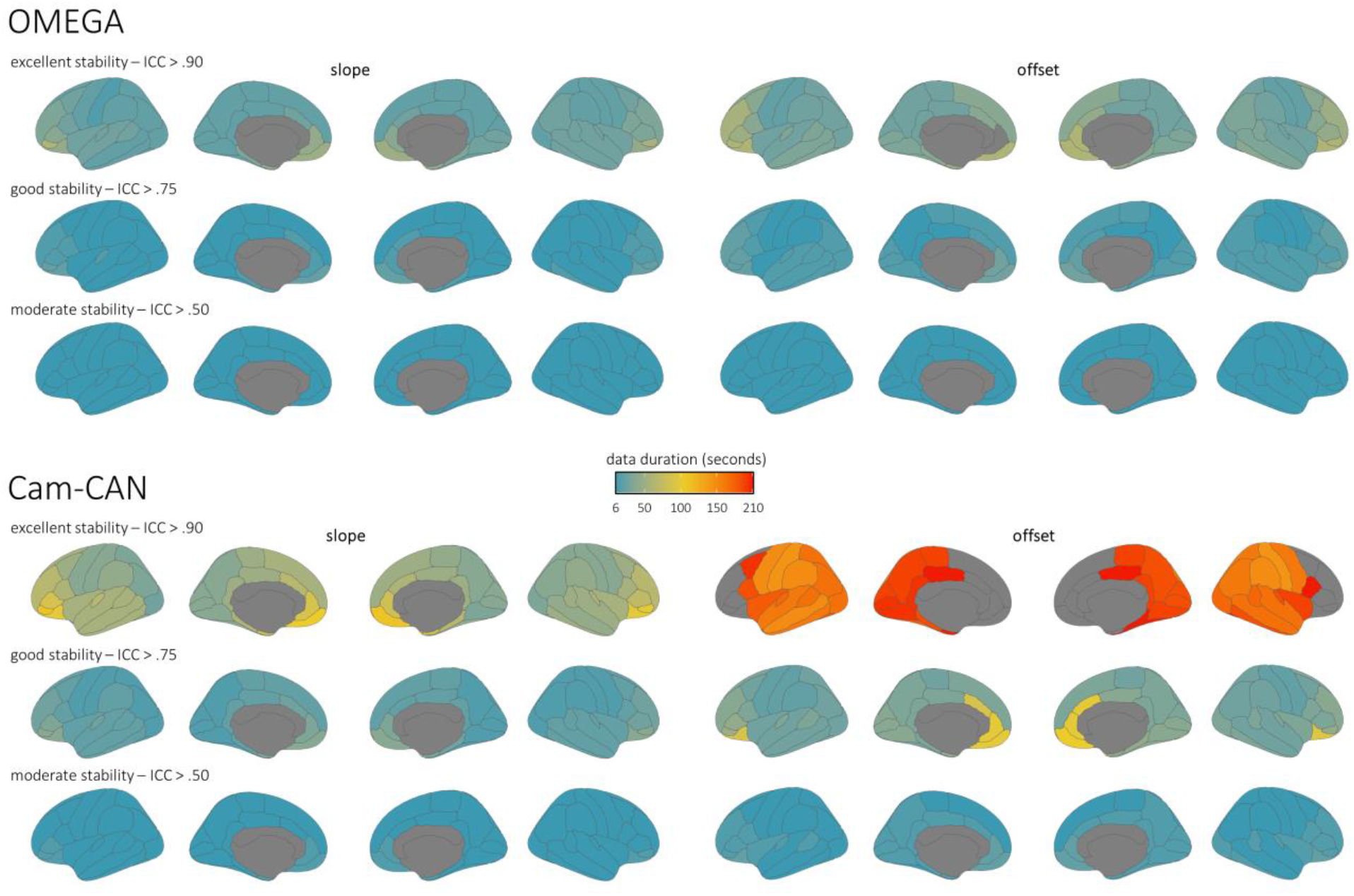
Brain maps of intra-session stability of parameterized aperiodic features. Parcellated surface maps per each aperiodic feature (left: slope; right: offset) indicate the length of data (color bar; in seconds) required to achieve accepted thresholds for moderate (ICC > .50; bottom rows), good (ICC > .75; middle rows), and excellent (ICC > .90; top rows) reliability in each participant sample (top: OMEGA, N = 107; bottom: Cam-CAN, N = 50) for each modeled cortical region of the Desikan-Killiany atlas. Warmer colors indicate worse temporal stability, and regions left grey did not achieve the indicated level of reliability within the maximum length of data available for the respective participant sample (OMEGA: 120 seconds; Cam-CAN: 210 seconds).

**Figure 5.**
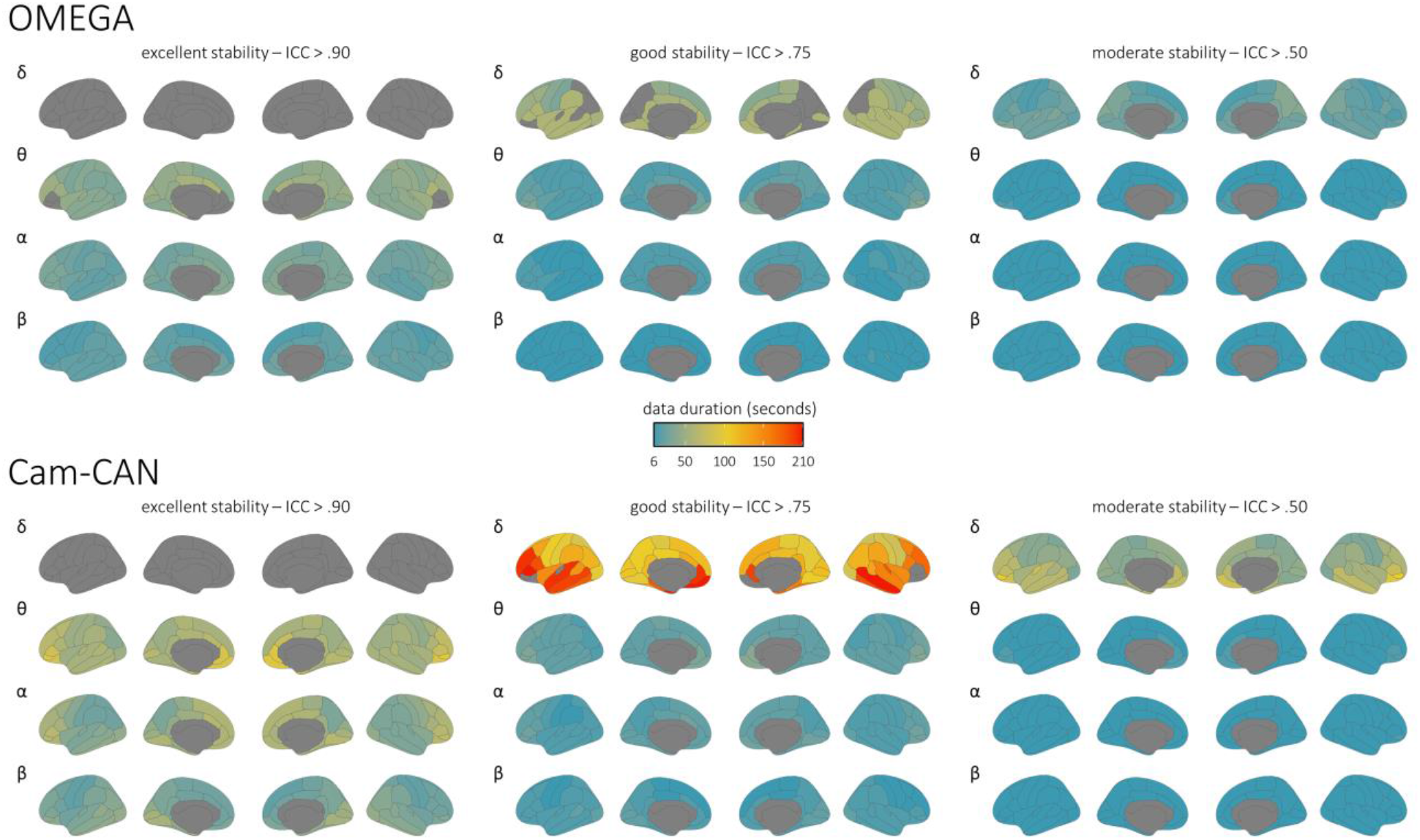
Brain maps of intra-session stability of parameterized periodic features. Parcellated surface maps per each spectral frequency (denoted by Greek letters to the top-left of each set) indicate the length of data (color bar; in seconds) required to achieve accepted thresholds for moderate (ICC > .50; right), good (ICC > .75; middle), and excellent (ICC > .90; left) reliability in each participant sample (top: OMEGA, N = 107; bottom: Cam-CAN, N = 50) for each modeled cortical region of the Desikan-Killiany atlas. Warmer colors indicate worse temporal stability, and regions left grey did not achieve the indicated level of reliability within the maximum length of data available for the respective participant sample (OMEGA: 120 seconds; Cam-CAN: 210 seconds).

### 3.3 Influence of demographics & cognitive factors

We investigated whether common participant sample characteristics impacted the stability of MEG spectral features. Even without correcting for multiple comparisons (6 frequencies × 4 sample characteristics = 24 tests), none of the effects were significant at *p* < .05. Post-hoc Bayesian testing indicated evidence for the null hypothesis in all cases except one (high-gamma & handedness; BF_01_ = 0.75; error % = 7.09 e^−6^; Figure 6).

**Figure 6.**
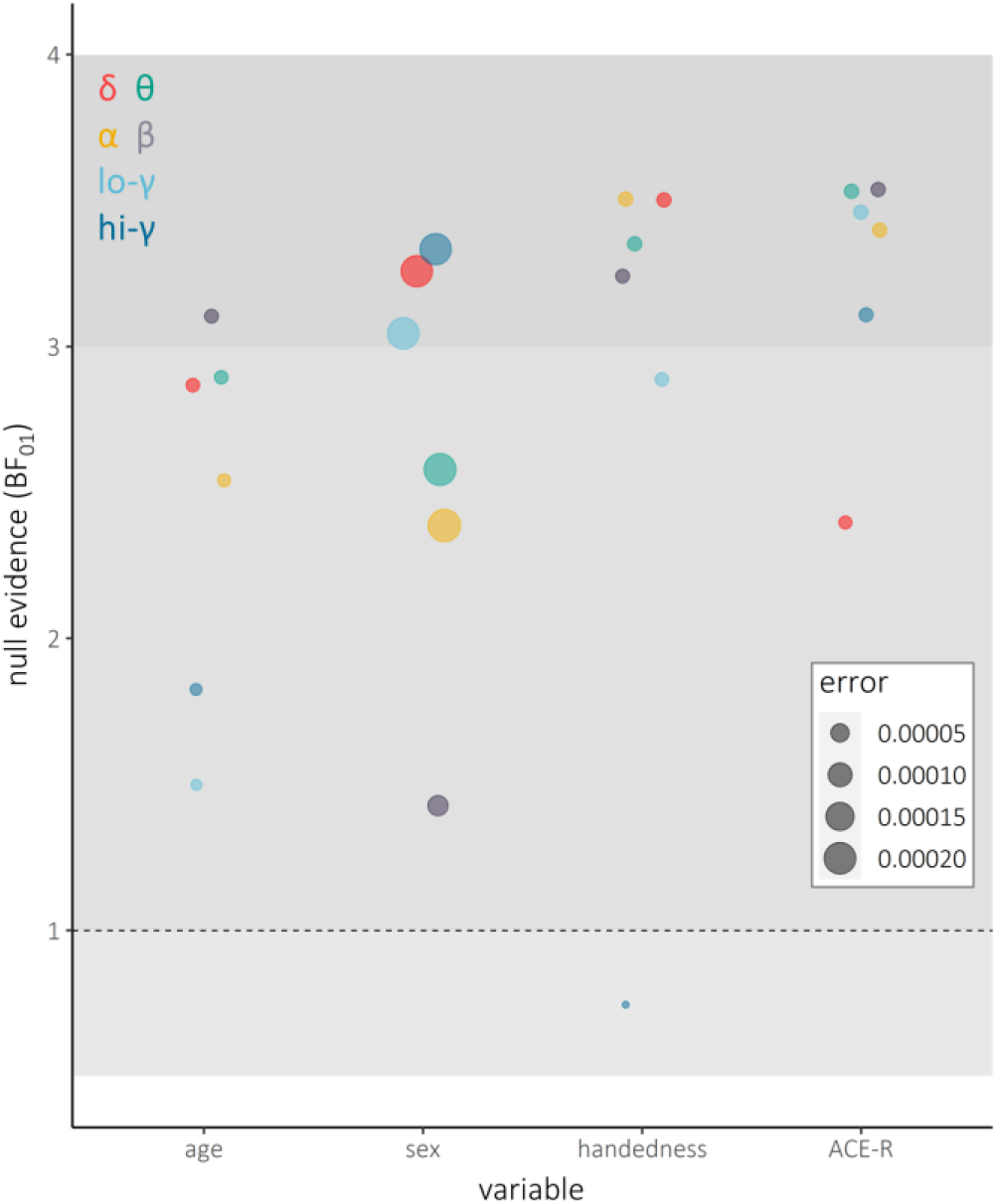
Bayesian model evidence for null effects of participant sample characteristics on the stability of spectral power estimates. Dots indicate Bayesian model evidence (y-axis; in BF01) and associated model errors (denoted by relative dot size) for null effects of common participant sample characteristics (x-axis) on temporal stability for each spectral frequency (denoted by dot color). Horizontal shaded intervals represent accepted thresholds for weak evidence for H1 (BF01 = .3 – 1), weak evidence for H0 (BF01 = 1 – 3), and moderate evidence for H0 (BF01 ≥ 3). Note that these models were only obtained for the Cam-CAN sample, due to its even distribution of age and availability of ACE-R cognitive scores.

## 4. Discussion

Despite a substantial literature examining the relationships between frequency-defined neural activity and various states of cognition and disease, little research exists regarding appropriate recording durations to derive robust spectral estimates of human electrophysiology signals. We believe that evidence-based recommendations concerning the minimum data duration required for extracting meaningful spectral features of brain activity are required to guide the design of resting-state studies and inform decisions when growing shared data repositories in the field. This work partially addresses this knowledge gap by quantitatively estimating the intra-session stability over time of source-imaged spectral features of resting-state MEG data.

Overall, we find that spectral estimates of task-free MEG brain activity stabilize remarkably quickly when derived from durations as short as 30 seconds, and typically 120 seconds, of data. These results align well with a recent study reporting that participants can be differentiated from one another based on MEG spectral “fingerprints” obtained with as little as 30 seconds of resting-state data (da Silva Castanheira et al., 2021). We noted some exceptions in the high-gamma range over frontal and temporal cortices, where the data duration required to reach stability was longer than for the alpha and beta bands. This indicates that, in cases where shorter MEG recording times are desirable, such as when collecting data from special or patient populations, the length of resting-state recordings could be determined depending on the cortical regions and spectral frequencies of interest. For instance, recordings of at least 2 minutes would be appropriate for investigations focused on ongoing alpha and beta activity, while studies of ongoing delta/theta/low-gamma activity would require at least 3 minutes of resting-state MEG, and those targeting high-frequency gamma power should be based on more than 3 minutes of contiguous data. We emphasize, however, that spectral instability at higher frequencies should not be conflated to signals being affected by intractable levels of noise. Indeed, the variability of high-frequency gamma activity is well documented to be associated to various cognitive functions (Başar, 2013; Uhlhaas et al., 2011; Ward, 2003; Wiesman et al., 2020; Wiesman and Wilson, 2019), disease states (Herrmann and Demiralp, 2005; Mably and Colgin, 2018; Uhlhaas and Singer, 2010), and inter-individual differences (da Silva Castanheira et al., 2021; Hirschmann et al., 2020; Muthukumaraswamy et al., 2010; Perry et al., 2013; Shaw et al., 2017). Furthermore, we show that estimates of intra-session resting-state MEG stability in normative populations require samples of at least 50 participants. These recommendations are based on our sub-sampling approach and offer researchers a yardstick when examining stability of spectral power derivatives at the group level.

We also found that the stability of spectral features was similar between both MEG instruments tested (CTF vs. MEGIN/Elekta), sampling rates used in data collection (2400 Hz vs. 1000 Hz), and resting-state paradigms (i.e., eyes-open vs. eyes-closed). We acknowledge the present study was not exhaustive with regards to all possible experimental factors that might influence MEG signal stability, but the MEG instruments and paradigmatic approaches encompass those most employed in the field. The similarity of results from data collected with different sampling rates also suggests that it is the absolute temporal duration (e.g., in seconds), rather than the number of data samples, that dictates the stability of spectral MEG features, at least when sampling at or above 1000 Hz. Additionally, the stability of spectral features was unaffected by common participant characteristics such as age, sex, handedness, and cognitive status. We emphasize that these factors are of research interest but evidence from our results indicate they may not require different durations of signal acquisition in the resting-state of healthy participants.

We also examined the robustness across varying epoch lengths of estimates of aperiodic and periodic spectral components of neurophysiological resting-state time series. The estimates of periodic components stabilized (ICC > .90) when derived from less than two minutes of data in the alpha and beta bands across the brain, and within 3.5 minutes of data in the theta band. We note that the theta, alpha, and beta bands are the most susceptible to exhibiting separable peaks in the human neurophysiological spectrum (Donoghue et al., 2020b). This contrasts with parameterized delta activity estimates, which did not reach excellent temporal stability in any region with less than 3.5 minutes of data. This lack of robustness was not explained by the short epoch duration and window length used for PSD derivations: using longer epochs (12 s) and window lengths (6 s) produced equivalent results. Rather, we found that the instability of low-frequency spectral estimates was due to variability of FOOOF models across participants. This indicates that the FOOOF algorithm is challenged in accounting for the periodic components in the delta band because it may be too close to the lower edge of the frequency spectrum, and/or that delta rhythmic activity is uncommon in healthy participants and therefore that estimates of periodic activity may be spurious. This latter point is aligned with the small number of delta peaks in resting-state data reported in the original FOOOF study (Donoghue et al., 2020b). Overall, low-frequency estimates of parameterized periodic spectral components in resting-state MEG are highly variable over time within typical session durations and should be interpreted with caution.

We found that estimates of the aperiodic slope were robust when derived from at least two minutes of OMEGA and Cam-CAN data. In contrast, the stability of the aperiodic offset differed between the OMEGA and Cam-CAN samples. Whereas the estimation of the spectral offset was stable for OMEGA data durations of about two minutes, its estimation from Cam-CAN data required longer durations and offset estimates in several frontal and medial regions did not reach excellent stability. We are unable to empirically determine the origin of this discrepancy in this study, as many experimental parameters (e.g., MEG instrument, resting-state paradigm and noise-cancellation approach) were different between the OMEGA and Cam-CAN samples. However, we can speculate that the parameters that most likely influence the low-frequency offset are the noise-cancellation method (i.e., hardware third-order synthetic gradiometers versus software MaxFilter spatial filter) and the resting-state paradigm (i.e., eyes-open vs. eyes-closed). The estimation of the aperiodic offset is highly dependent on the correct estimation of lower-frequency periodic components, namely from the delta band, in which stability was also challenged in these cortical regions. Future research is needed to explore the sources of this disagreement between the OMEGA and Cam-CAN samples.

We also found that the estimation of high-gamma spectral power was not stable over the recording durations available from OMEGA and Cam-CAN. We can extrapolate from our data that future studies will require MEG resting-state recordings of at least 10 minutes per participant to derive stable estimates of high-gamma power (Figure S1). This pattern of high intra-session gamma variability and low intra-session alpha and beta variability is similar to a recent study of long term (i.e., 3 year) inter-session variability of MEG spectral power (Lew et al., 2021). This suggests that alpha and beta spectral power represent a reliable estimate of inter-participant neural variability, both within a single-session “snapshot” of participant neural activity and across sessions separated even by multiple years. We are also aware that our findings may not generalize to all populations. The variability of brain activity may be different in e.g., younger children or specific patient populations for reasons either related to participant compliance with recording conditions or actual differences in neurophysiological activity due to development of pathology.

Considering these limitations, we look at our present findings as a normative benchmark for neuroscience studies of resting-state brain activity. We hope that our recommendations will prove particularly useful to researchers when designing their MEG resting-state studies and extracting spectral features. Amid concerns about the reproducibility and replicability of findings across scientific fields, it may be reassuring that common estimators of spectral properties of brain signals are robust when derived from recording durations typically used in a large majority of existing research and open datasets.

## 5. Acknowledgements

This work was supported by a grant F32-NS119375 (AIW) from the National Institutes of Health and a doctoral fellowship from NSERC (JDSC). Data collection and sharing for this project was provided by the Open MEG Archives (OMEGA) and the Cambridge Centre for Ageing and Neuroscience (Cam-CAN). Cam-CAN funding was provided by the UK Biotechnology and Biological Sciences Research Council (grant number BB/H008217/1), together with support from the UK Medical Research Council and University of Cambridge, UK. OMEGA and the Brainstorm app are supported by funding to SB from the NIH (R01-EB026299), a Discovery grant from the Natural Science and Engineering Research Council of Canada (436355-13), the CIHR Canada research Chair in Neural Dynamics of Brain Systems, the Brain Canada Foundation with support from Health Canada, and the Innovative Ideas program from the Canada First Research Excellence Fund, awarded to McGill University for the Healthy Brains for Healthy Lives initiative. The funders had no role in study design, data collection and analysis, decision to publish, or preparation of the manuscript.

## Supplementary Figures

**Figure S1.**
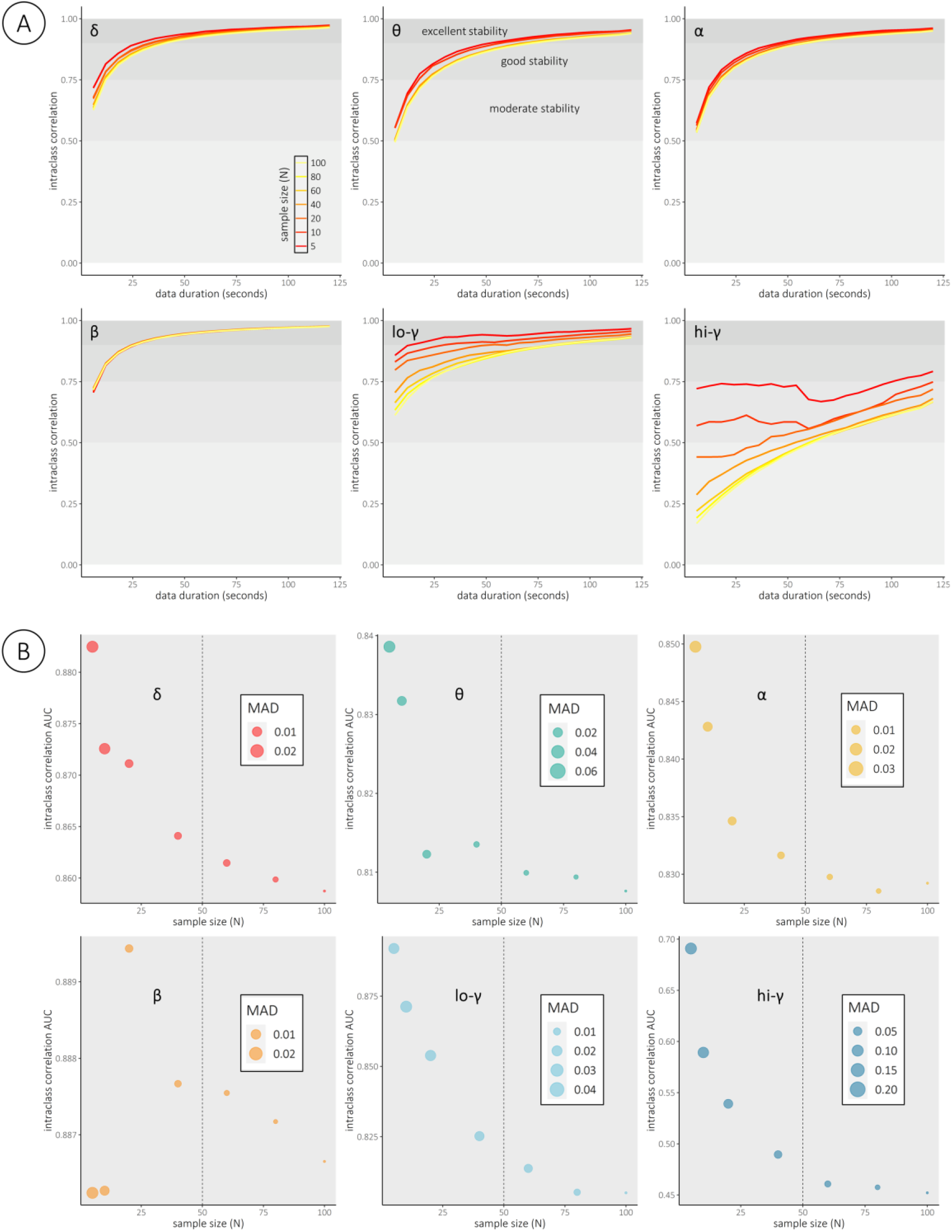
Intra-session temporal stability time series per sample size and spectral frequency. (A) Intraclass correlation coefficients (y-axes) are medians over 1,000 permutations of epoch order, and represent the stability of each neural power estimate, averaged over all cortical regions, as a function of participant sample size (denoted by line color; median over 100 permutations of participants), length of data (x-axes; in seconds), and spectral frequency (separate plots; denoted by Greek letters in the top left of each). Horizontal shaded intervals in each plot represent accepted thresholds for moderate (ICC > .50), good (ICC > .75), and excellent (ICC > .90) reliability. (B) Intraclass correlation area-under-the-curve (AUC) values are integrals of the curves in (A), and dot sizes represent integrals for similar median absolute deviation (MAD) curves. The y-axes in (B) vary across each frequency sub-plot to accentuate effects of sample size. Note that this modeling was only performed in the OMEGA sample, to quasi-empirically estimate the number of participants that would subsequently be needed from the Cam-CAN dataset.

**Figure S2.**
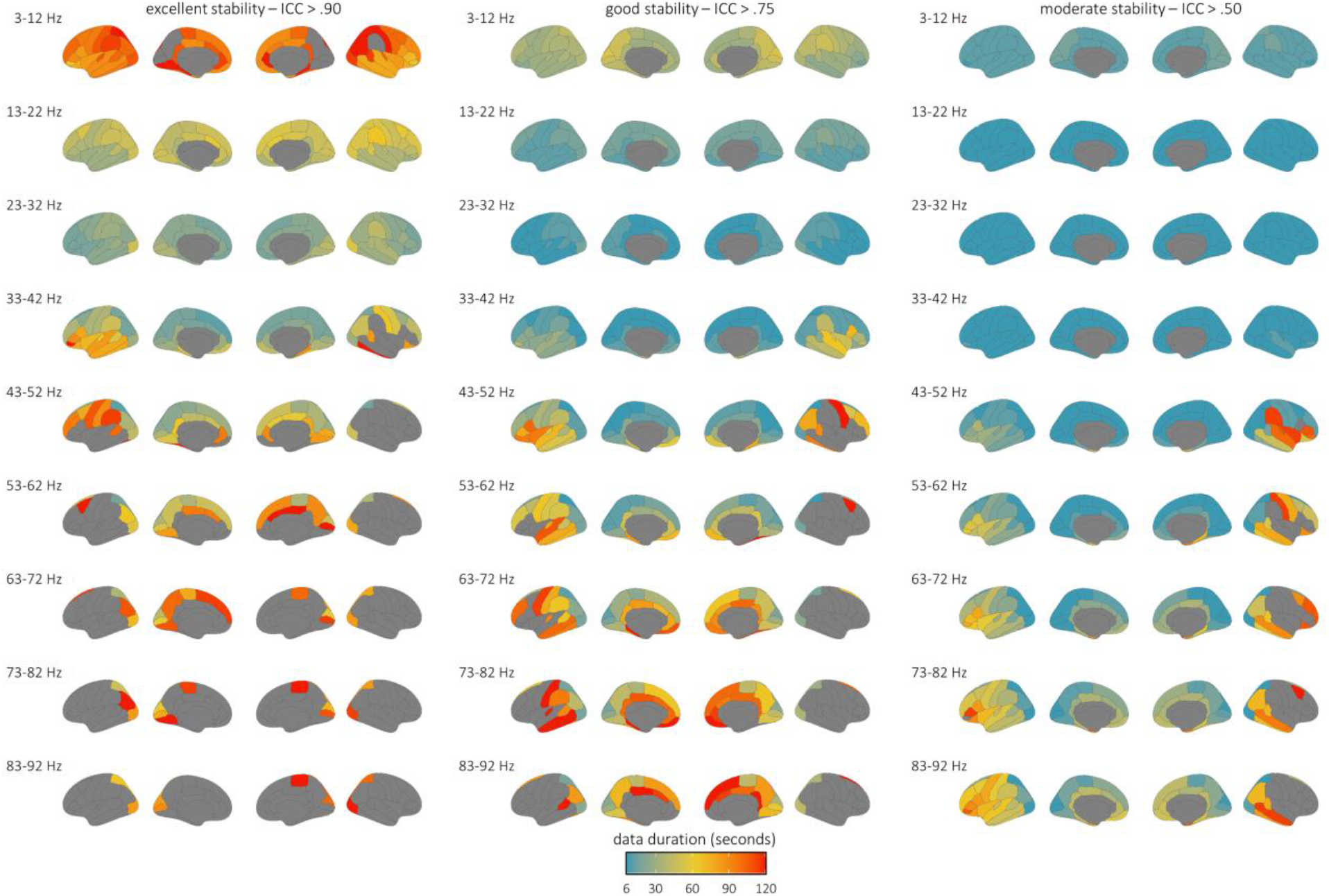
Regional intra-session temporal stability per participant sample and 10 Hz spectral band. Parcellated surface maps are equivalent to the OMEGA surface maps in Figure 3, but computed on data averaged over equivalently-sized 10 Hz spectral definitions (denoted by the text to the top-left of each set), rather than canonical frequency bands.

**Figure S3.**
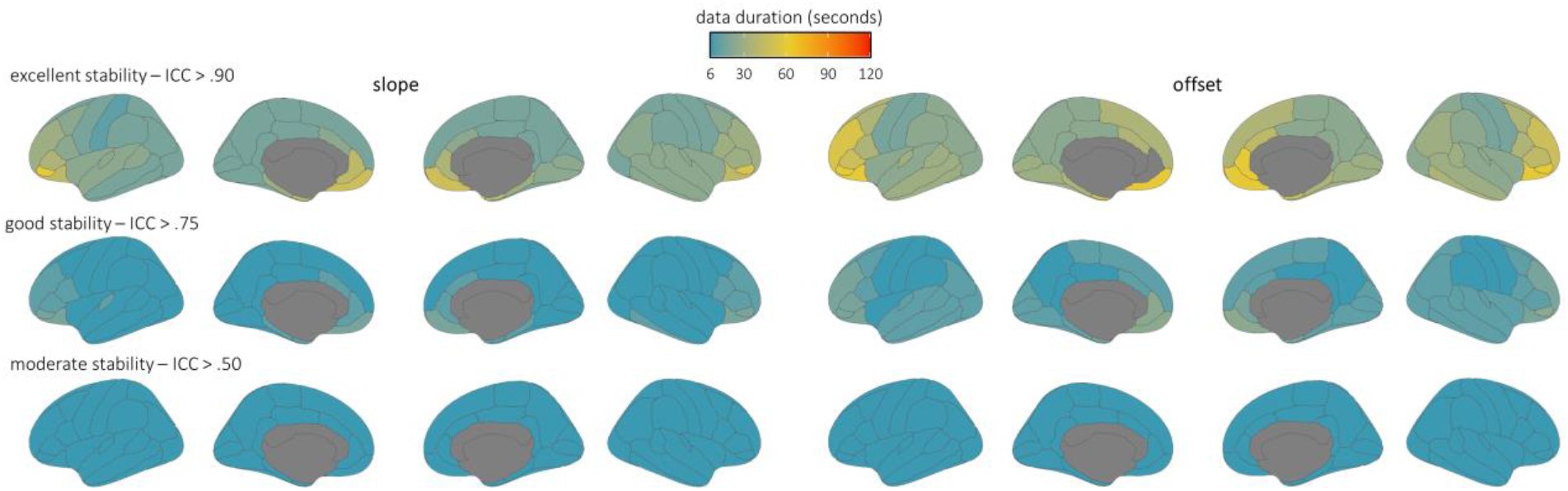
Regional intra-session temporal stability of parameterized aperiodic features using 12 seconds epochs. Parcellated surface maps are equivalent to the OMEGA surface maps in Figure 4, but computed using longer epochs and PSD windows (original: epoch = 6 seconds, PSD window = 3 seconds; new: epoch = 12 seconds; PSD window = 6 seconds).

**Figure S4.**
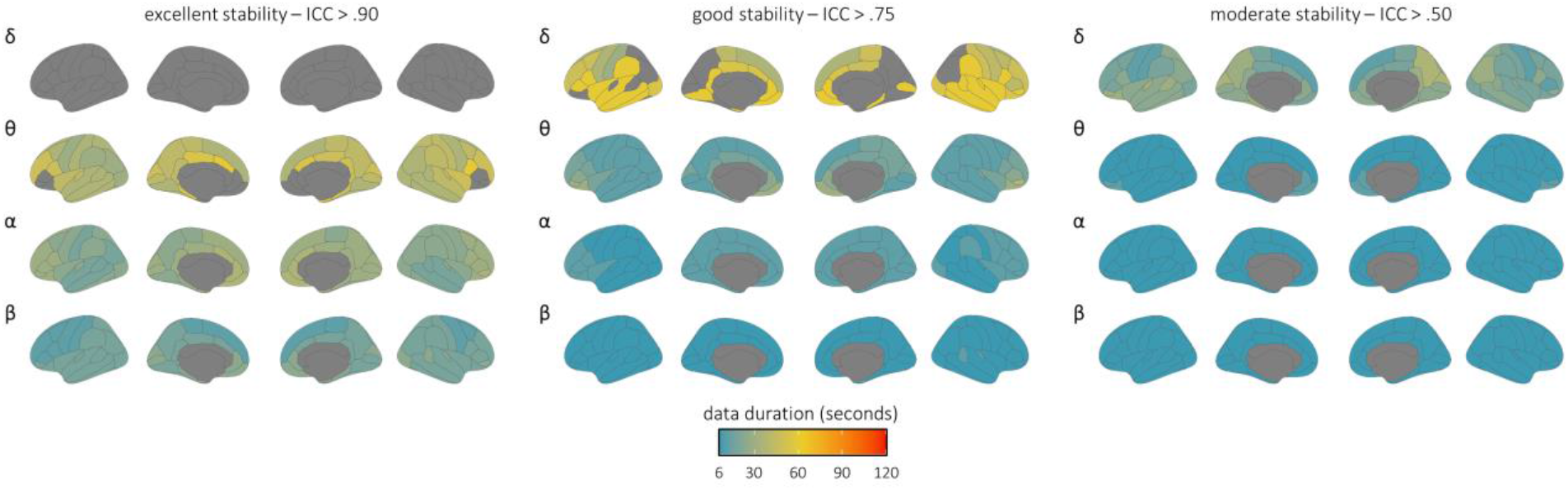
Regional intra-session temporal stability of parameterized periodic features using 12 seconds epochs. Parcellated surface maps are equivalent to the OMEGA surface maps in Figure 5, but computed using longer epochs and PSD windows (original: epoch = 6 seconds, PSD window = 3 seconds; new: epoch = 12 seconds; PSD window = 6 seconds).

**Figure S5.**
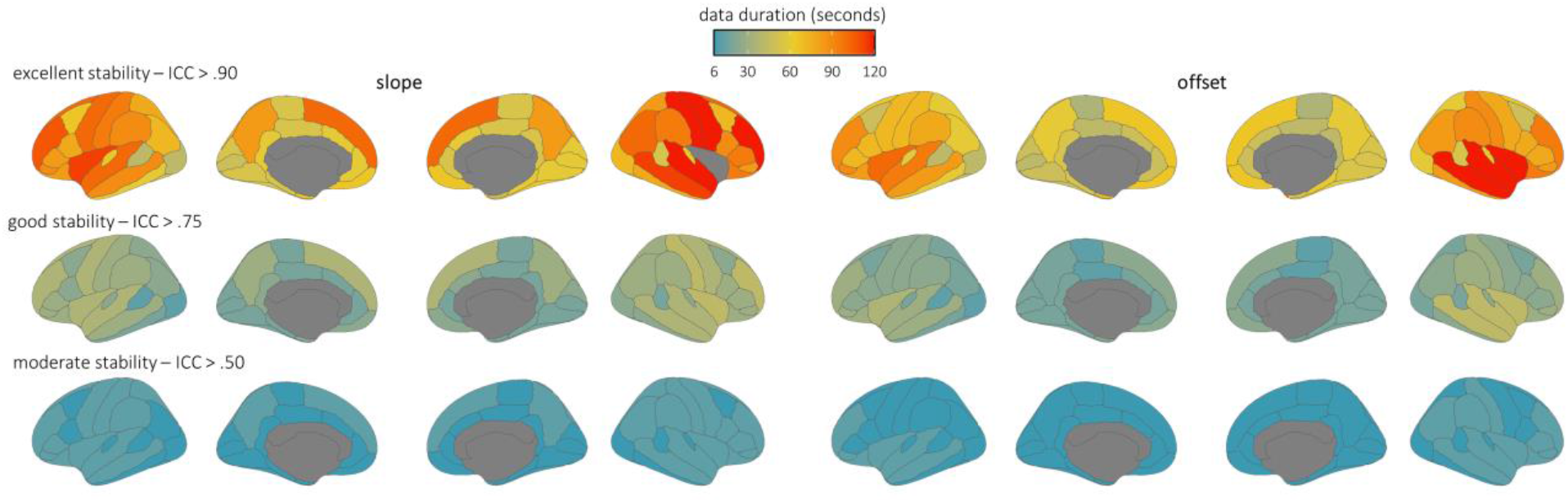
Regional intra-session temporal stability of parameterized aperiodic features covarying FOOOF model fit. Parcellated surface maps are equivalent to the OMEGA surface maps in Figure 4, but computed on the residuals from regressions of parameter estimates on FOOOF model fit (R2) across participants, per each cortical region.

**Figure S6.**
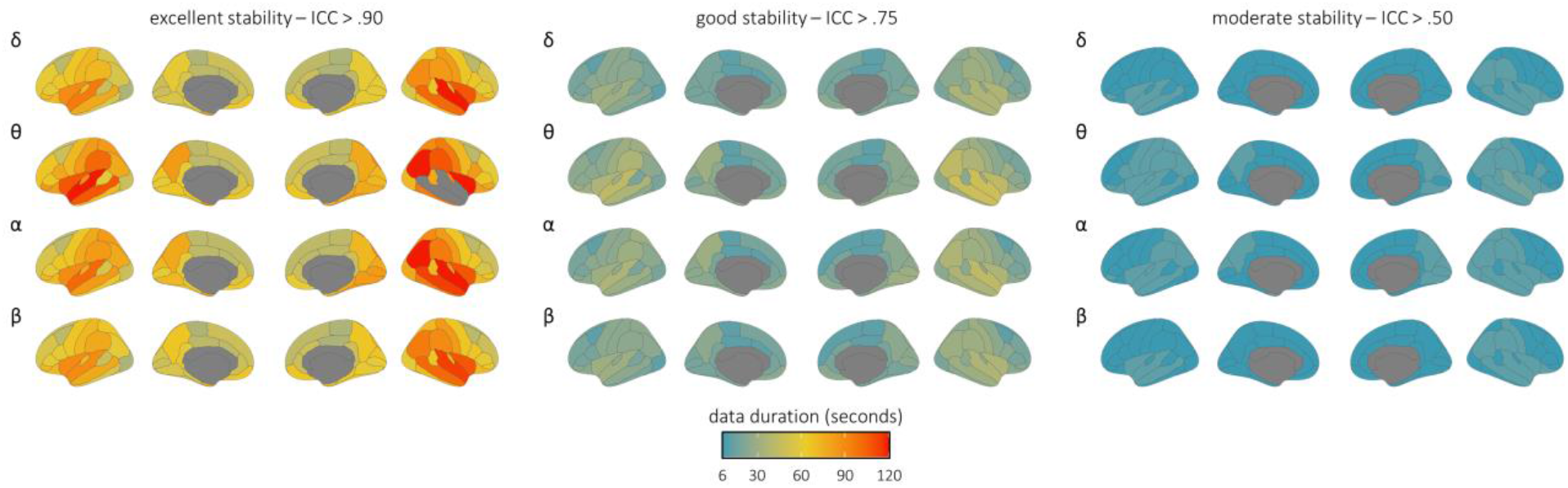
Regional intra-session temporal stability of parameterized periodic features covarying FOOOF model fit. Parcellated surface maps are equivalent to the OMEGA surface maps in Figure 5, but computed on the residuals from regressions of parameter estimates on FOOOF model fit (R2) across participants, per each cortical region.

